# Active outer hair cell motility can suppress vibrations in the organ of Corti

**DOI:** 10.1101/2020.01.03.893933

**Authors:** T. Jabeen, J. C. Holt, J. R. Becker, J.-H. Nam

## Abstract

High sensitivity and selectivity of hearing require active cochlea. The cochlear sensory epithelium, the organ of Corti, vibrates due to external and internal excitations. The external stimulation is acoustic pressures mediated by the scala fluids, while the internal excitation is generated by a type of sensory receptor cells (the outer hair cells) in response to the acoustical vibrations. The outer hair cells are cellular actuators that are responsible for cochlear amplification. The organ of Corti is highly structured for transmitting vibrations originating from acoustic pressure and active outer hair cell force to the inner hair cells that synapse on afferent nerves. Understanding how the organ of Corti vibrates due to acoustic pressure and outer hair cell force is critical for explaining cochlear function. In this study, excised cochlear turns were freshly isolated from young gerbils. The organ of Corti in the excised cochlea was subjected to mechanical and electrical stimulation that are analogous to acoustical and cellular stimulation in the natural cochlea. Organ of Corti vibrations including those of individual outer hair cells were measured using optical coherence tomography. Respective vibration patterns due to mechanical and electrical stimulation were characterized. Interactions between the two vibration patterns were investigated by applying the two forms of stimulation simultaneously. Our results show that the interactions could be either constructive or destructive, which implies that the outer hair cells can either amplify or suppress vibrations in the organ of Corti. We discuss a potential consequence of the two interaction modes for cochlear frequency tuning.

**Statement of Significance:** The function of the mammalian cochlea is characterized by sharp tuning and high-level of amplification. Both tuning and amplification are achieved mechanically through the action of cellular actuators in the sensory epithelium. According to widely accepted theory, cochlear tuning is achieved by ‘selectively amplifying’ acoustic vibrations. This study presents a set of data suggesting that the cochlear actuators can both amplify and suppress vibrations to enhance cochlear tuning. Presented results will explain why the actuator cells in the cochlea spend energy in the locations where there is no need for amplification.

## Introduction

The cochlea encodes sounds into neural signals according to frequency and amplitude. As sound waves enter the cochlea through the middle ear bones, inter-scala pressure waves propagate from the base to the apex of the cochlea (1). The traveling waves peak at different locations such that low frequency waves propagate farther toward the apex than high frequency waves. As a result, a specific location of the cochlea responds best to a specific frequency, called the tonotopy. Cochlear tonotopy is closely related to the mechanical gradient along the cochlear length, which is dominated by the basilar membrane (2, 3). The auditory epithelium, called the organ of Corti (OoC), sits on the basilar membrane and is covered by the tectorial membrane to form the OoC complex. Relative motion (shear) between the OoC and the tectorial membranes results in activation/inhibition of the mechano-receptive cells— the inner and the outer hair cells (4). The outer hair cells elongate or shrink as their transmembrane potential changes (5). Vibrations of the OoC complex are then modulated by mechanical feedback from the outer hair cells so that small sounds at best-frequency are amplified (6). As such, the outer hair cells contribute to the key functions of the cochlea—tuning and amplification of sounds.

OoC vibrations are more complicated than it was thought prior to the finding of outer hair cell electromotility. Classical understanding of OoC mechanotransduction is based on kinematics, treating the OoC as a rigid body (7–9). The classical model explains how the stereocilia are deflected by the relative motion between the OoC and the tectorial membrane. This mechanism of stereocilia deflection underlies the foundation of theoretical models (10–14). The rigid body approximation, however, is being relaxed as more experimental data are revealing complex relative motions among OoC fine structures. For example, the top and the bottom surfaces of the OoC (often represented by the reticular lamina and the basilar membrane) have been shown to vibrate with considerable phase difference. This relative motion among OoC structures is dependent on the level and frequency of stimulation (15, 16). Outer hair cell electromotility affects the reticular lamina more than the basilar membrane (17–20). While there is relatively good agreement that outer hair cell electromotility is essential for cochlear amplification and tuning, how those functions are achieved remains controversial. For example, some argue that cochlear amplification is achieved by longitudinal accumulation of energy (18, 21, 22), while others consider that cochlear amplification is essentially local (20). While the outer hair cells are considered to provide power for cochlear amplification, some researchers argue that the outer hair cells rather attenuate intense sounds (23).

The subject of this study is the modulation of OoC vibration patterns due to outer hair cell motility. We used excised cochlear tissues from young gerbils to investigate how OoC vibrations due to mechanical and electrical stimulation interact. We captured the deformation of individual outer hair cells using optical coherence tomography (OCT). Our results demonstrate that vibration patterns generated by the two stimuli can either constructively or destructively interact. The consequences of these interactions are discussed.

## Methods

### Tissue preparation

Cochleas harvested from young Mongolian gerbils (two to five weeks old) were used for experiments according to the institutional guidelines of the University Committee on Animal Resources (UCAR) at the University of Rochester. Gerbils were deeply anesthetized with isoflurane after which the cochlea was acutely isolated and placed in a petri dish containing: in mM, 145 Na-Gluconate, 7 NaCl, 3 KCl, 5 NaH_2_PO_4_, 0.1 MgCl_2_, 5 D-Glucose, 0.1 CaCl_2_, 5 HEPES. The acidity and the osmolarity of solutions used in this study were adjusted to pH 7.35 and 300 mOsm, respectively. In the petri dish, the cochlea was further reduced to a single cochlear turn. This study targeted 1-mm, mid-to-apical sections centered at 8 mm from the basal end (red colored section in Fig. 1A). Basal and apical turns were removed using forceps and sharp blades. Bones between scalae were removed from both basal and apical sides to expose the cochlear epithelium from the section of interest (Fig. 1B). Apical and basal openings of the remaining cochlear coil were sealed using cyanoacrylate glue (orange colored in Fig. 1C and D). The reduced cochlear turn was then transferred to a microfluidic chamber initially containing the same perilymph-like solution. Cyanoacrylate sealant was applied along the circumference of the cochlear turn (Fig. 1C). Tissue preparation typical required 60 to 80 minutes.

**Figure 1.**
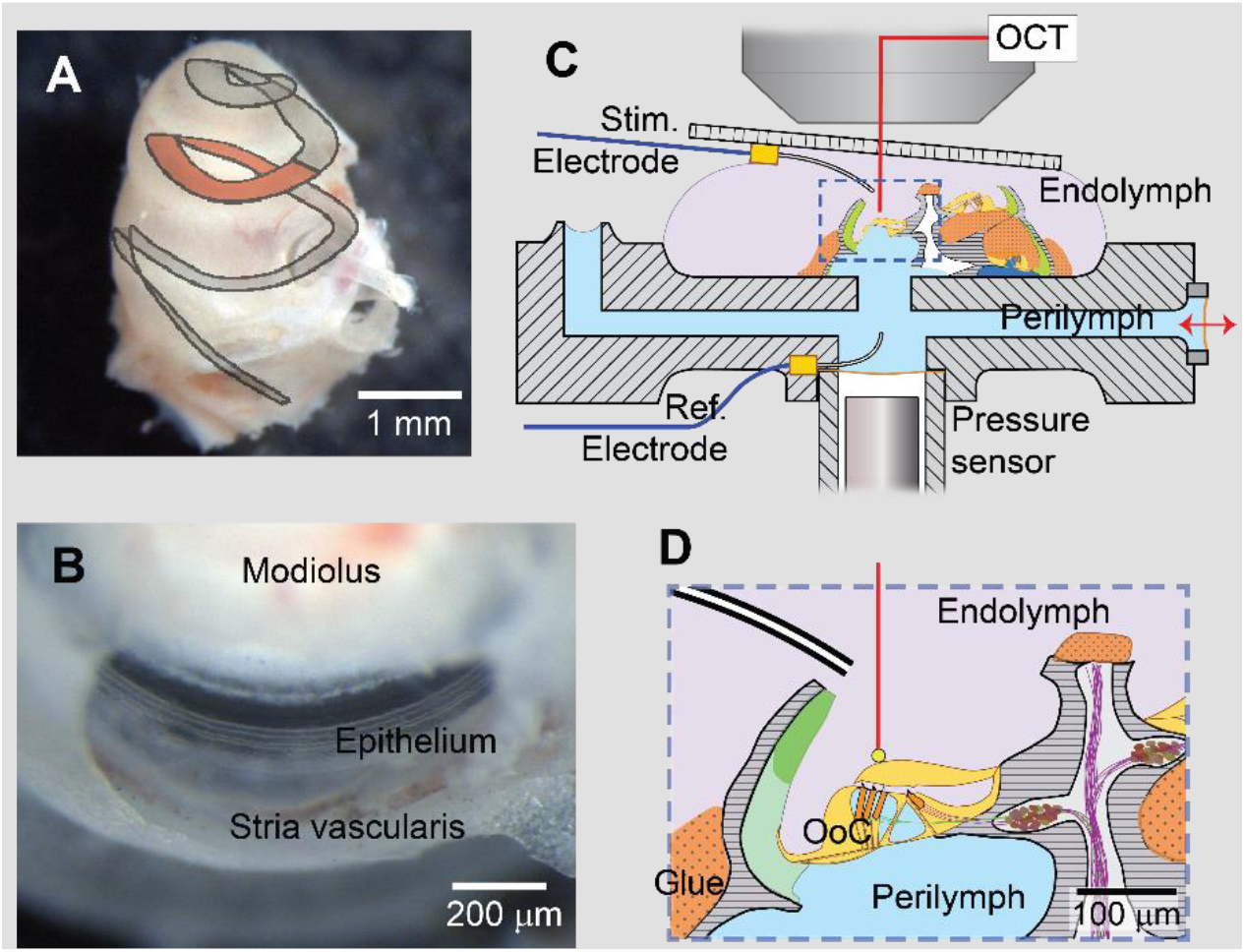
Preparation of excised cochlear turn. (**A**) Excised gerbil cochlea. The apical and the basal turns are removed to expose the target location (red section). (**B**) Excised cochlear turn placed in microfluidic chamber. (**C**) Schematic of prepared tissue in the microfluidic chamber. The excised cochlear turn is sealed with glue to separate two fluid spaces. The tissue is stimulated mechanically through a stimulating port (red arrow) and/or electrically through a pair of electrodes. Resulting vibrations are measured using optical coherence tomography. (**D**) An enlarged view of the organ of Corti (OoC) in the chamber from Panel C.

### Microfluidic chamber

Our microfluidic chamber was fabricated using a stereolithography printer (Moai, Peopoly). Mechanical stimulation was delivered by vibrating an opening covered with elastic membrane with a piezoelectric actuator (red arrow in Fig. 1C). Another opening was provided for pressure release. There was a pair of inlet-outlet ports to refresh the solution in the chamber (not shown). Electrical stimulation across the OoC was applied through a pair of electrodes (Fig. 1C). After the cochlear tissue was attached and sealed to the chamber, bottom and top fluids were replaced with an artificial endolymph (145 KCl, 0.1 CaCl_2_, 4 HEDTA, 10 K-HEPES, 8 Glucose, 2 Na-Pyruvate, concentrations in mM) and a second perilymph (145 Na-Gluconate, 7 NaCl, 3 KCl, 5 NaH_2_PO_4_, 0.1 MgCl_2_, 5 D-Glucose, 1 CaCl_2_, 5 HEPES, concentrations in mM), respectively.

### Stimulation and measurements

Transepithelial electrical stimulation was applied through a pair of Ag/AgCl electrodes. Alternating current (typically between 50 and 200 µA) was supplied by a current source (CS580, Stanford Research Systems), or a custom-built current generator. The voltage across the tissue was also monitored. Typical electrical impedance between two electrodes was 4-6 kΩ. Mechanical stimuli were delivered using a piezoelectric actuator (PC4WM, Thorlabs, NJ) driven by a high-voltage amplifier (E-505.00, Physik Instrumente). The PZT actuator tip vibrated with an amplitude of 30 nm per 0.5 V driving voltage. At 1 kHz, this stimulation level generated about 1 Pa of pressure beneath the slit. Stimulus functions were generated using Matlab code that controls a data acquisition board (PCI-6353, National Instruments). For vibration measurements, a commercial optical coherence tomography (OCT) imaging system (Ganymede, Thorlab) was used. The system uses a light source with 900 nm center wavelength, and its scanning unit has an A-scan rate of 100 kHz. The system was modified to use a 20x objective (N.A. 0.4, Mitutoyo) to enhance optical resolution. The imaging system was driven with a Matlab program to operate its M-scan mode for vibration measurements. For validation of OCT vibration measurements, a piezoelectric actuator was vibrated under the OCT system and then under a laser interferometry system (MMA 300 system, Polytec). Measurement results from the two vibrometry systems were compared. The difference between two measurement systems was less than 5 percent within 0.1 to 20 kHz frequency range. Between 0.3 and 3 kHz, the root-mean-square noise of OCT vibration measurements was 0.5 nm. Most experiment protocols ran a set of measurements either at different scanning points (spatial sweep), at different frequencies (frequency sweep), at different phases between stimulations (phase sweep), etc. To avoid potential artifacts due to measurement sequence, the order of events was randomized.

### Viability of tissue

Per the purpose of this study, the viability of prepared tissue is most explicitly represented with the level of outer hair cell electromotility. All presented data were obtained when the outer hair cell electromotility was greater than 100 nm/mA. Besides electromotility, two morphological features served as good indicators for the state of each preparation. First, the tectorial membrane attachment to the OoC was particularly vulnerable to surgical or chemical insults. Therefore, a secure intact attachment was a good indicator for a preparation’s viability. Second, outer hair cell shape was sensitive to incomplete separation between the two fluid compartments. In these instances, OoC deformation due to swelling outer hair cells was evident within 10 minutes of artificial endolymph application. We declared a proper seal when leakage between the two fluid-spaces was < 0.16 μL/min at slit pressures of 20 Pa (Marnell *et al*., 2018). When these two conditions were positive at early stages, the electromotility stayed near or above 10 nm per 100 µA current for over 180 minutes post cochlear isolation. All presented data were obtained when: 1) the tectorial membrane was attached to the OoC; 2) leakage between two fluid spaces was negligible; and 3) outer hair cell electromotility was above 10 nm per 100 µA current.

## Results

### Fine OCT imaging helps to discern tissue status

Our *ex vivo* preparation combined with OCT provided high resolution imaging of OoC micro structures. The image signal was stronger for the reticular lamina, outer hair cells, pillar cells, Hensen’s cells, basilar membrane fiber layers, and the bottom layer of tectorial membrane, while it was weaker in Deiters’ cells and the body of the tectorial membrane (Fig. 2A). Micrometer-level anatomical characteristics could be effectively resolved including the gaps between the three rows of outer hair cells, the sub-tectorial space and nerve fibers passing through the tunnel of Corti. Such resolution enabled us to observe the deformation of individual outer hair cells throughout the experiment. These images were helpful in monitoring the status of the preparation, especially when the tissue was deformed due to stress or damage. For example, as outer hair cells swelled, they shrank along their length, and the reticular lamina deflected downward, while the Hensen’s cell outside the third-row outer hair cell bulged upward. As this deformation progressed, the tectorial membrane became detached from the OoC at its lateral extreme. When outer hair cells had smooth texture and distinct boundaries, their motility was good. In contrast, a granular texture and blurry boundaries of the outer hair cells were associated with a loss in electromotility. The micro-mechanics seemed local: Even if a radial section showed swollen outer hair cells with little electromotility, the outer hair cells in other sections of the same preparation could be in good shape with strong electromotility. In our preparation, the basilar membrane was tilted between 20 and 30 degrees from the horizontal line (Fig. 2A). In this orientation, the outer hair cells were roughly aligned to the optical axis which was helpful in measuring their length change (electromotility).

**Figure 2.**
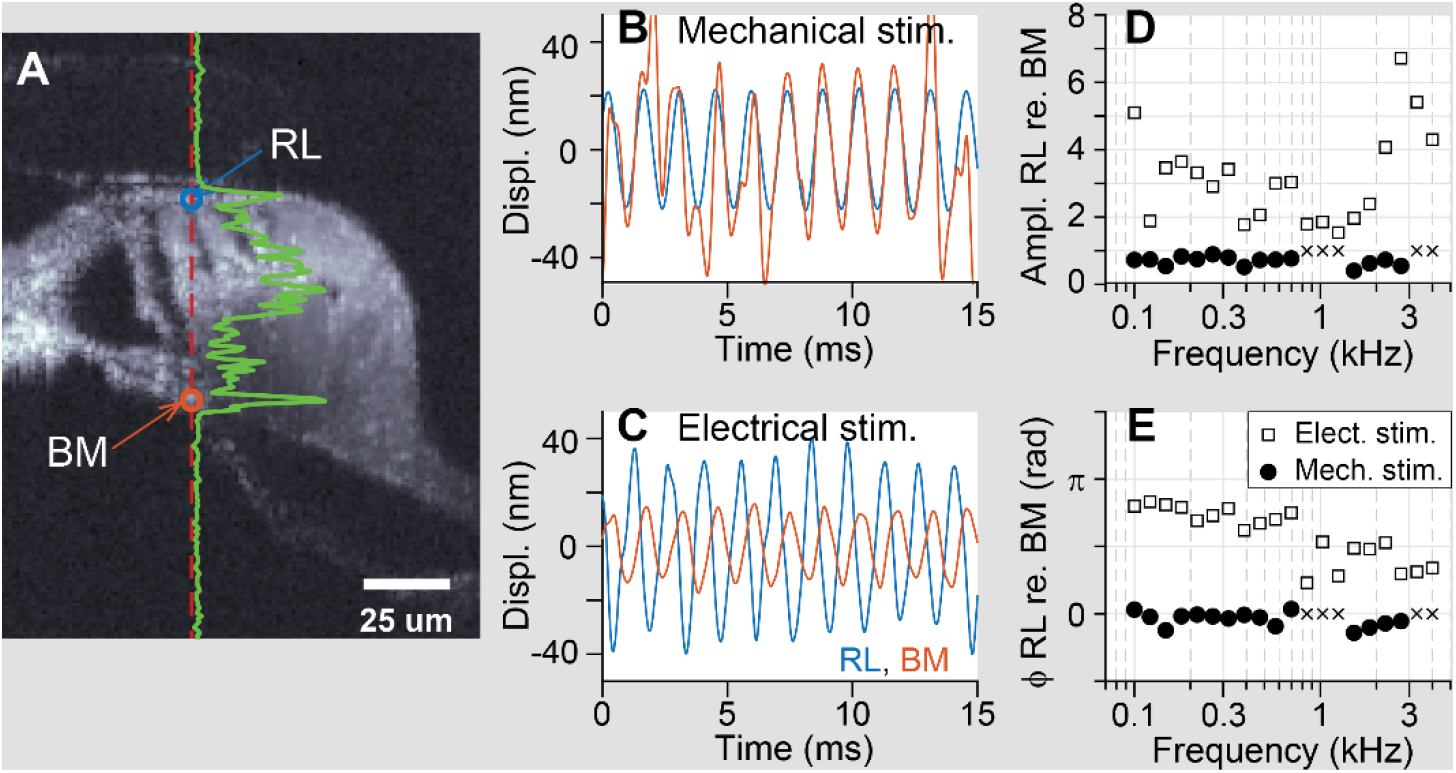
OoC vibrations due to mechanical and electrical stimulation. (**A**) B-scan image of the OoC. Red vertical dashed line represents the optical axis where vibrations were measured. The horizontal distance between the curve from the vertical line (green curve) indicates signal strength along the optical depth. Two void circles indicate the measurement points presented in panels (B-E). (**B**) Vibrations of the reticular lamina (blue) and the basilar membrane (red) due to mechanical stimulation. (**C**) Vibrations due to electrical stimulation (100 µA at 1 kHz). (**D**, **E**) Relative motion (amplitude and phase) of the reticular lamina with respect to the basilar membrane at different stimulating frequencies. Measurement location was 9 mm from the basal end. The cross symbols (×) indicate data points with poor signal where signal-to-noise ratio < 8 dB.

### Two vibration patterns

Vibration measurements were made from regions 7 to 9 mm from the basal end of the gerbil cochlea, which corresponds to the frequency range of 1-3 kHz. Note that our excised preparation compromises natural mechanical and electrical boundary conditions including the pressure delivery along the length of the scalae. Therefore, instead of examining the relationship between input stimulation and tissue vibration, we analyzed the relative motion between micro-structures within the OoC. Similar to other studies (15, 18, 24), the reticular lamina and the basilar membrane were chosen for analysis, because they represent the top and bottom of the OoC, and because the optical signal was stronger at these locations. We elected to use a calcium level of 0.1 mM in the upper chamber (scala media space), which is higher than reported physiological levels. This choice was a compromise between two considerations: to decrease the mechano-transduction current with higher than physiological calcium-level (25) so that we could better separate the mechanical and electrical responses, and to maintain the shape of the tectorial membrane with a calcium-level not too far from physiological conditions (26).

Vibration patterns due to mechanical and electrical stimulations were distinctly different (Fig. 2). When mechanically stimulated, the reticular lamina and the basilar membrane vibrated in phase and the vibrations of the reticular lamina were smaller than those of the basilar membrane (Fig. 2B). This trend regarding the relative motion between the top and bottom of the OoC was highly consistent across different stimulating frequencies (symbol ● in Fig. 2D and E). The ratio in vibration amplitude between the top and bottom of the OoC (between ‘RL’ and ‘BM’ in Fig. 2A) was between 0.5 and 1. Our observations of passive mechanics are in line with the classical kinematic model (7) in that the ratio between the top and bottom of the OoC is approximately 1. Our measurements also agree with reported ratios between the reticular lamina and the basilar membrane made in other *in vivo* observations (16, 24 Chen, Nuttall, 2011). When electrically stimulated, however, the reticular lamina vibrated at a different phase from the basilar membrane (> 60 degrees, Fig. 2E). The relative displacement between the top and bottom of the OoC depended on stimulating frequency (symbol □ in Fig. 2D and E).

Spatial patterns of OoC vibration were also different in response to the two types of stimulation (Fig. 3). These plots were obtained from 50 and 40 scan lines across the radial span of the OoC for mechanical- and electrical-stimulation cases, respectively, corresponding to a spacing of 3.0 and 3.6 µm between scan lines. For each scan, a sinusoidal stimulation (20 ms ramp time and 100 cycles at plateau) was applied and the resulting vibrations were measured. Data pixels with signal-to-noise ratios > 8 dB are represented with colored dots. Mechanical vibration patterns show that the vibration amplitude is greater in the middle of the OoC and decreases toward the lateral edges (Fig. 3A). This pattern reflects the anatomy where ligaments hold the medial and lateral ends of the basilar membrane. The peak displacement (the brightest spot in the amplitude plot) occurred near the joint between arcuate and pectinate zones of the basilar membrane and at the center of the OoC near the outer hair cells, in agreement with previous observations (17, 27). When the focal plane was at the level of the outer hair cells, the tectorial membrane did not have strong optical signals, but occasionally its top or bottom surface produced signals. We did not observe a phase difference between the tectorial membrane and the reticular lamina.

**Figure 3.**
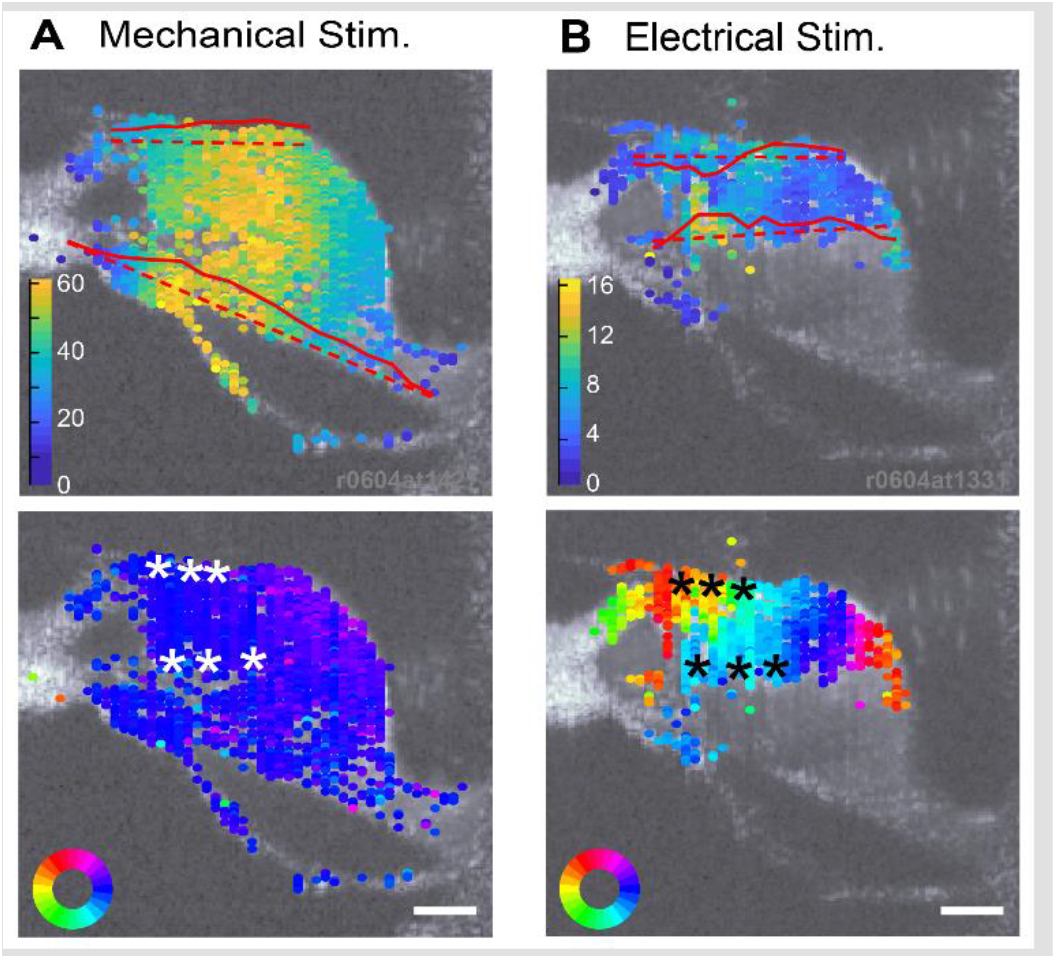
Two vibration patterns in the OoC. The amplitude and phase measurements of 50 and 40 A-scan lines are shown on top of corresponding B-scan image. The top and bottom row panels show amplitude and phase of vibrations, respectively. (**A**) Amplitude and phase of vibrations due to mechanical stimulation. The stimulating port was vibrated at 1 kHz with a 30 nm amplitude. (**B**) Amplitude and phase of vibrations due to electrical stimulation. Stimulating current was applied at 2 kHz with a 100 µA amplitude. **Top panels**: The red curves indicate vibrating shapes along the broken lines. Color bar units are nm. **Bottom panels**: Asterisk symbols indicate the extremities of three outer hair cells. The color rings at the left bottom corner indicate the color scale of phase angle. *E.g*., between the dark blue and yellow spots, there is a phase difference of 180 degrees. The scale bars indicate 25 μm.

Image resolution was fine enough to resolve the motility of individual outer hair cells. For the case shown in Fig. 3B, the phase difference between the top and bottom of the three rows of outer hair cells (asterisk symbols) were 162, 149, 60 degrees. With a current amplitude of 100 μA, the amplitude of electromotility was 28, 22 and 8 nm for the first-, second- and third-row cells, respectively. This trend of greater motility for the first- and second-row cells versus the third-row cell was observed in majority of cases (Fig. 4A and B). Despite such observations, we cannot exclude potential experimental artifacts such as faster deterioration of the third-row outer hair cells.

**Figure 4.**
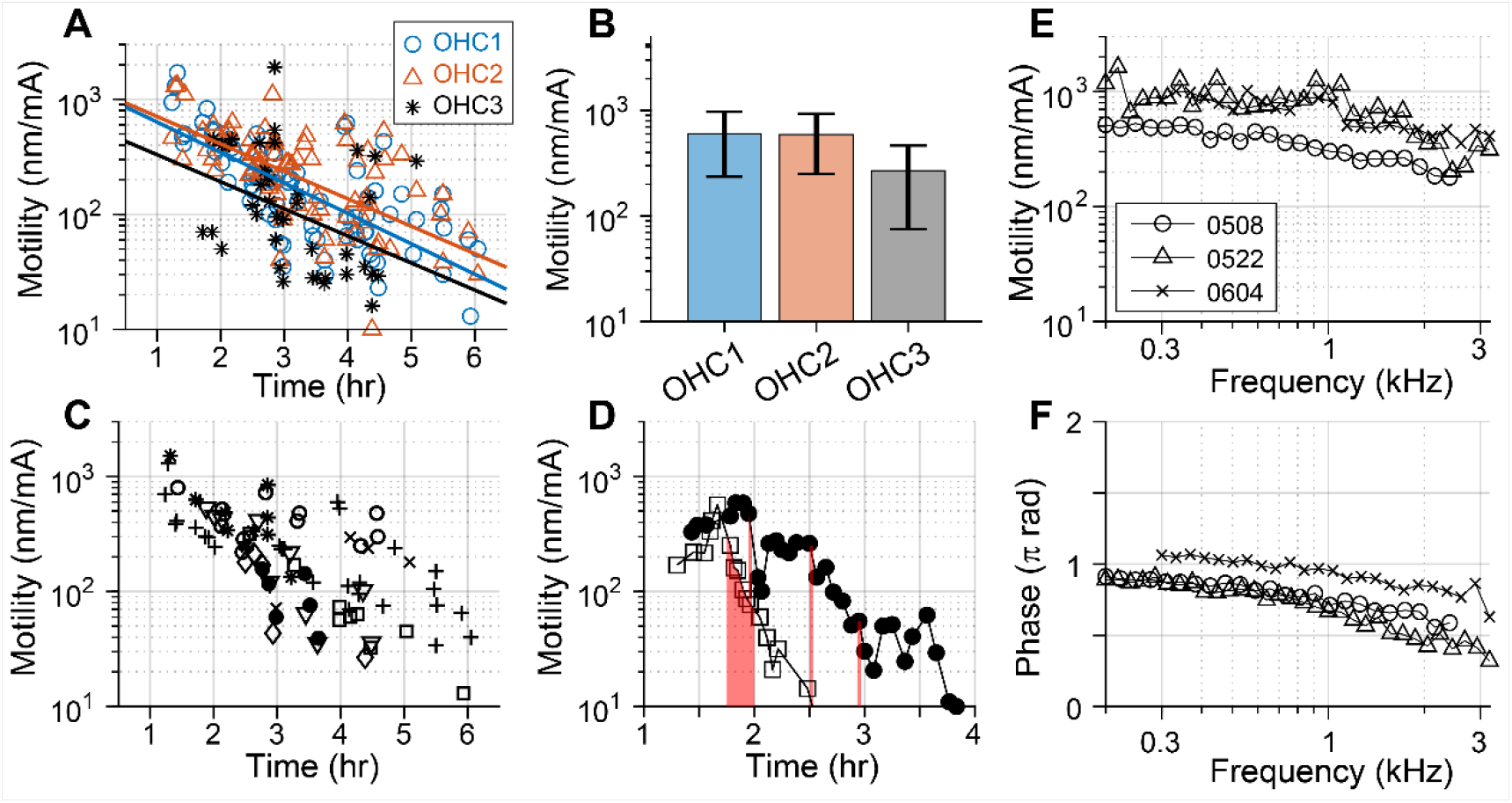
Electromotility of individual outer hair cells. Electromotility of outer hair cells was measured over time and frequency. Time zero is defined as the onset of experimental dissection. (**A**) Motility of individual cells from eight cochleae are presented. The lines indicate the trend of exponential decay for the three rows of outer hair cells distinguished by different symbols and colors (see legends). (**B**) Mean motility of the three rows of outer hair cells over the first hour of measurement (< 2.25 hr). The error bar indicate standard deviation. (**C**) Motility from different cochleae are distinguished by different symbols. (**D**) Na-Salicylate decreased outer hair cell electromotility. Shaded vertical columns indicate the span of Na-Salicylate application. The different symbols indicate independent trials. (**E, F**) Amplitude and phase of motility w.r.t electrical stimulation at different stimulating frequencies. Data are from three different cochleae.

### Outer hair cell’s electromotility

The electromotility of outer hair cells was defined as a length change in the cell body per transepithelial current. The electrical resistance between endolymphatic and perilymphatic electrodes was about 5 kΩ. The first measurements of individual preparations were made typically 1.25 hours after cochlear isolation. The time-course of outer hair cell electromotility decay varied across preparations (Fig. 4A). This variation is ascribed to different factors including the extent of surgical insults, chemical conditions, different measurement locations, and level of stimulation. In most cases, motility was greater than 100 nm/mA for 3 hours or longer. Larger and longer electrical stimulations tended to exhaust the preparation quicker, while modest level of mechanical stimulation (peak displacement at the reticular lamina < 100 nm) did not appear to affect tissue conditions. The motility of individual hair cells is presented in Fig. 4A. When analyzed for early hour groups (within one hour from measurement onset), the third-row outer hair cells had smaller motility compared to the other rows (Fig. 4B). This difference may be attributed in part to the trend for quicker swelling of the third-row outer hair cell. Considering the trend of motility decay over time (lines in Fig. 4A), the electromotility of outer hair cells from our measurement location (8 mm from the basal end of the gerbil cochlea) is expected to be greater than 1000 nm/mA. In Fig. 4C, each data point represents the average motility of measurable outer hair cells within observed radial section (mostly outer hair cell row 1 or 2). Na-salicylate, a blocker of outer hair cell electromotility (28), reduced the motility (Fig. 4D). In a case with 15 minutes of exposure to 10 mM Na-salicylate (□), the motility decreased quickly (>90 percent drop within an hour) and did not recover. In another trial (●), the tissue sample was subjected to Na-salicylate at three 2-minute exposures with 30-minute intervals. After the first application the motility reduced by 80 % and then recovered to the half of the pre-treatment motility. In subsequent applications, however, both inhibition and recovery were smaller. The outer hair cell’s electromotility decreased by less than an order of magnitude as stimulating frequency increased from 200 to 3200 Hz (Fig. 4E and F). The phase of motility, defined as the phase of elongation with respect to top-to-bottom chamber current, decreased approximately by 90 degrees over the frequency range, from 180 to 90 degrees. That is, as the frequency increases, the motility changed from conductive to capacitive, which is in-line with the Evans-Dallos model (29). As we have not investigated other cochlear locations, it is unclear if this trend is true regardless of location. Our measurements are consistent with previously reported observations. Karavitaki and Mountain (22, 30) reported radial displacements of 400 and 2000 nm/mA at 120 Hz from the apical and middle turn locations of the gerbil cochlea. Note that we present outer hair cell elongation instead of reticular lamina displacement. The measurements in Zha *et al*.’s (31) are analogous to our measurements in that they estimated outer hair cell length changes from the differential motion between the reticular lamina and basilar membrane. In their study, outer hair cells in basal locations (Characteristic frequency, CF = 20 kHz) of the gerbil cochlea deformed up to a few nanometers. Regarding the Na-salicylate treatment, a previous study showed 50 % of reduction in outer hair cell motility after 5 minutes application of 10 mM Na-salicylate and the motility was fully recovered within minutes (32).

### Vibrations due to simultaneous stimulation

Mechanical and electrical stimulation were applied simultaneously to observe how vibrations due to different sources interact (Fig. 5). A modest level of stimulation was applied to minimize quick exhaustion of the tissue. The chosen response level (10-30 nm vibration amplitude at the reticular lamina) is within the physiological range (*e.g*., (33)), and well above the noise floor of our measurements. Three different phases between the mechanical and electrical stimulation were tested (ϕ_ME_ = 0, 90 and 180 degrees). Reflecting the phase variation in the horizontal direction of the electrically-stimulated case, the spatial pattern of vibration amplitude was affected by ϕ_ME_. Among the three measured cases (ϕ_ME_ = 0, 90, and 180 degrees, Fig. 5), when ϕ_ME_ = 0, the reticular lamina vibrated least. When ϕ_ME_ = 180 degrees, the reticular lamina vibrated greatest. Notably, the lateral wall of the OoC (the rectangles in Fig. 5A) vibrated differently from the rest of OoC.

**Figure 5.**
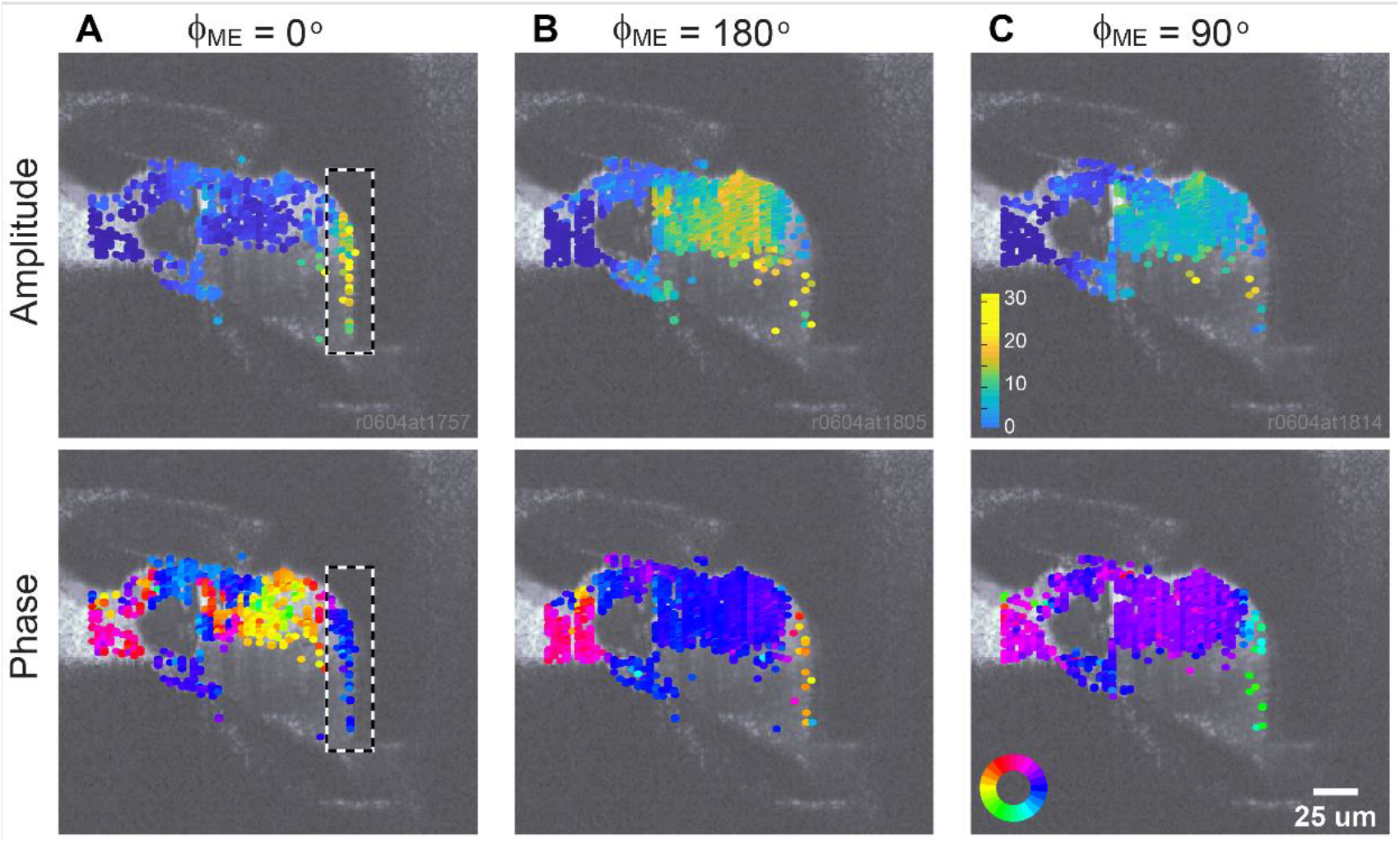
OoC vibration patterns due to simultaneous stimulations. Mechanical and electrical stimulations (1 kHz sinusoids) were applied simultaneously, but with different phases between the two stimulations (ϕ_ME_) so that electrical stimulation lags the mechanical stimulation by (**A**) zero degrees, (**B**) 180 degrees, and (**C**) 90 degrees. The upper and the lower row panels represent the vibrations in amplitude and phase, respectively. The scale bars shown only in the right column as the same scales were use in the other columns.

The interaction between vibrations due to the different stimulating methods was investigated further. The OoC vibrations were measured at 12 different phases between mechanical and electrical stimulations (ϕ_ME_ = 0, 30, 60, …, 360 degrees) for nine different frequencies between 0.3 and 1.5 kHz (Fig. 6). To decide on the level of the two stimulations, frequency responses due to mechanical or electrical stimulation alone were measured (columns A and B). When the stimulating port was vibrated with the same amplitude (mechanical stimulations), the OoC vibration peaked at 0.8 kHz, and the phase between the stimulation and the tissue vibration changed by 0.5-1 cycle over the test frequency range. For electrical stimulation, the reticular lamina response was relatively flat over the tested frequency range while the basilar membrane response decayed as the frequency increased. Then, mechanical and electrical stimulation were applied simultaneously. When one stimulation overwhelms the other, it is not ideal to observe interactions between the two vibration patterns. Therefore, we adjusted the levels of mechanical stimulation so that the reticula laminar vibrated 10 nm across the frequency range. When this equalized mechanical stimulation and the electrical stimulation were applied simultaneously, a vibration amplitude of ~20 nm was expected (10 nm from each stimulations).

**Figure 6.**
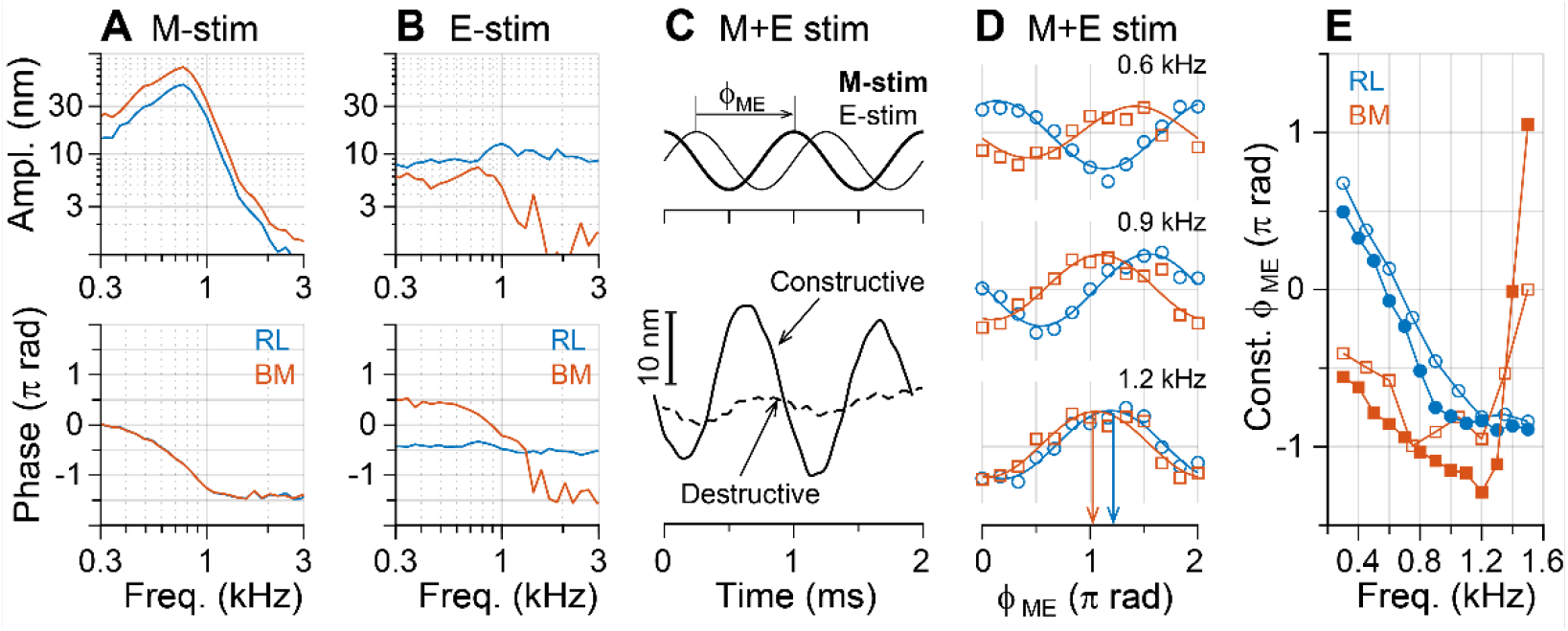
Constructive and destructive interactions between two vibrations. Measurements were made at two points of the OoC--the reticular lamina (RL) and the basilar membrane (BM). (**A**) Frequency response to mechanical stimulations. (**B**) Frequency response to electrical stimulations. (**C**) Top plot: Simultaneous stimulations. Electrical stimulation lags mechanical stimulation by ϕ_ME_. Bottom plot: Depending on ϕ_ME_, responses due to the two stimulations add up (constructive) or cancel each other (destructive). (**D**) Normalized vibration amplitude at the reticular lamina (RL, ○) and basilar membrane (BM, □) for different ϕ_ME_ values. Results at three frequencies are shown. The constructive ϕ_ME_ is defined at the peak of fitted sinusoidal curve (arrows). (**E**) Constructive ϕ_ME_ versus stimulating frequency for the reticular lamina and the basilar membrane. The filled and void symbols represent two sets of data obtained from different locations of a preparation.

There existed a specific value of ϕ_ME_ at which the resulting response due to simultaneous stimulations was constructive or destructive (add up, or cancel each other, Fig. 6C). As expected from the summation of two sinusoidal curves, the constructive ϕ_ME_ is 180 degrees apart from the destructive ϕ_ME_. The vibration amplitudes were obtained at different ϕ_ME_’s, while other conditions including measurement point, stimulating amplitudes and frequency remained the same. The data points of response amplitude versus ϕ_ME_ were fitted with a sinusoidal curve from which the constructive ϕ_ME_ was obtained (Fig. 6D). The constructive ϕ_ME_ varies over stimulating frequency (Fig. 6E). As was implicated in Figs. 5 and 3B, the interaction was not uniform across the OoC. For the case presented in Fig. 6, the reticular lamina and basilar membrane had similar constructive ϕ_ME_ near 1.2 kHz, or both the top and bottom surfaces of the OoC will constructively respond when stimulated with ϕ_ME_ ≈ −150 degrees at 1.2 kHz. On the contrary, at 0.3 kHz, constructive ϕ_ME_ of the reticular lamina and the basilar membrane are approximately 180 degrees apart. That is, at 0.3 kHz, when one side of the OoC is under constructive interaction, the other side of the OoC will be under destructive interaction.

Different modes of interaction between mechanically and electrically-evoked OoC vibrations can affect frequency selection at a cochlear location. An example of frequency selection through constructive and destructive interaction is demonstrated in Fig. 7. Two ϕ_ME_ values were chosen that were 180 degrees apart (−120 and 60 degrees). Due to interactions between the two stimulations, the vibration amplitude of the reticular lamina varied between 10 and 30 nm depending on frequency (top panel of Fig. 7A). Near 0.8 kHz, the constructive and destructive interaction contrasted most prominently. The difference between the two curves (ϕ_ME_ = −120 and 60 degrees) is reminiscent of a tuning curve (bottom panel of Fig. 7A). Consistent with the difference between the reticular lamina and the basilar membrane responses in Fig. 6B, the frequency responses measured at the two locations were different. That is, a condition that amplifies the reticular vibration could suppress the basilar membrane vibrations.

**Figure 7.**
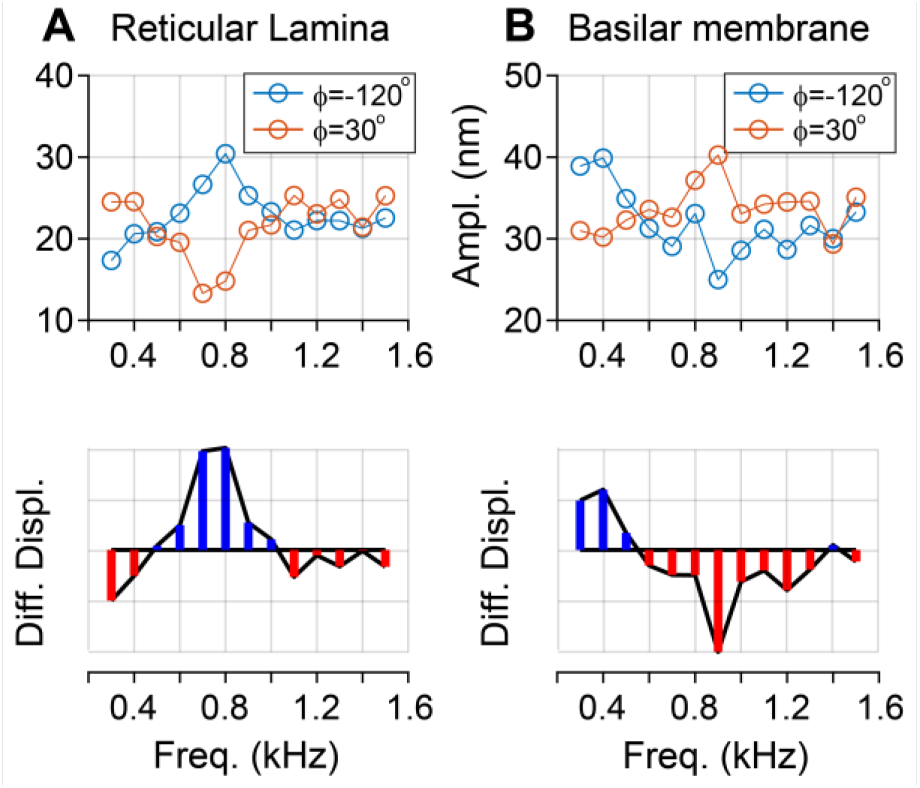
Synthesized frequency response. (**A**) Response of the reticular laminar to simultaneous stimulation. (**B**) Response of the basilar membrane to simultaneous stimulation. Mechanical stimulation level was adjusted to have a flat vibration amplitude of 10 nm at the reticular lamina.

## Discussion

Active motility of outer hair cell is required for both sensitivity and selectivity of hearing. Apparently, uniform amplification over frequency (Fig. 8C) will not enhance frequency tuning. Therefore, the prevailing view on cochlear tuning is that the active feedback from the outer hair cells *selectively* amplifies vibrations at the best frequency, specific to a location (34, 35). The mechanism of this selective amplification has been a central theme in hearing research. In theory, an actuator in a vibrating system could be used for either amplification or suppression, if the timing of actuation varies depending on frequency. The outer hair cells can operate more proactively for frequency tuning by using both amplification and suppression depending on frequency (Fig. 8D). In the intact cochlea, it is not trivial to manipulate or estimate the timing of actuation (*i.e*., outer hair cell motility) because passive and active responses are coupled together to form a feedback loop.

**Figure 8.**
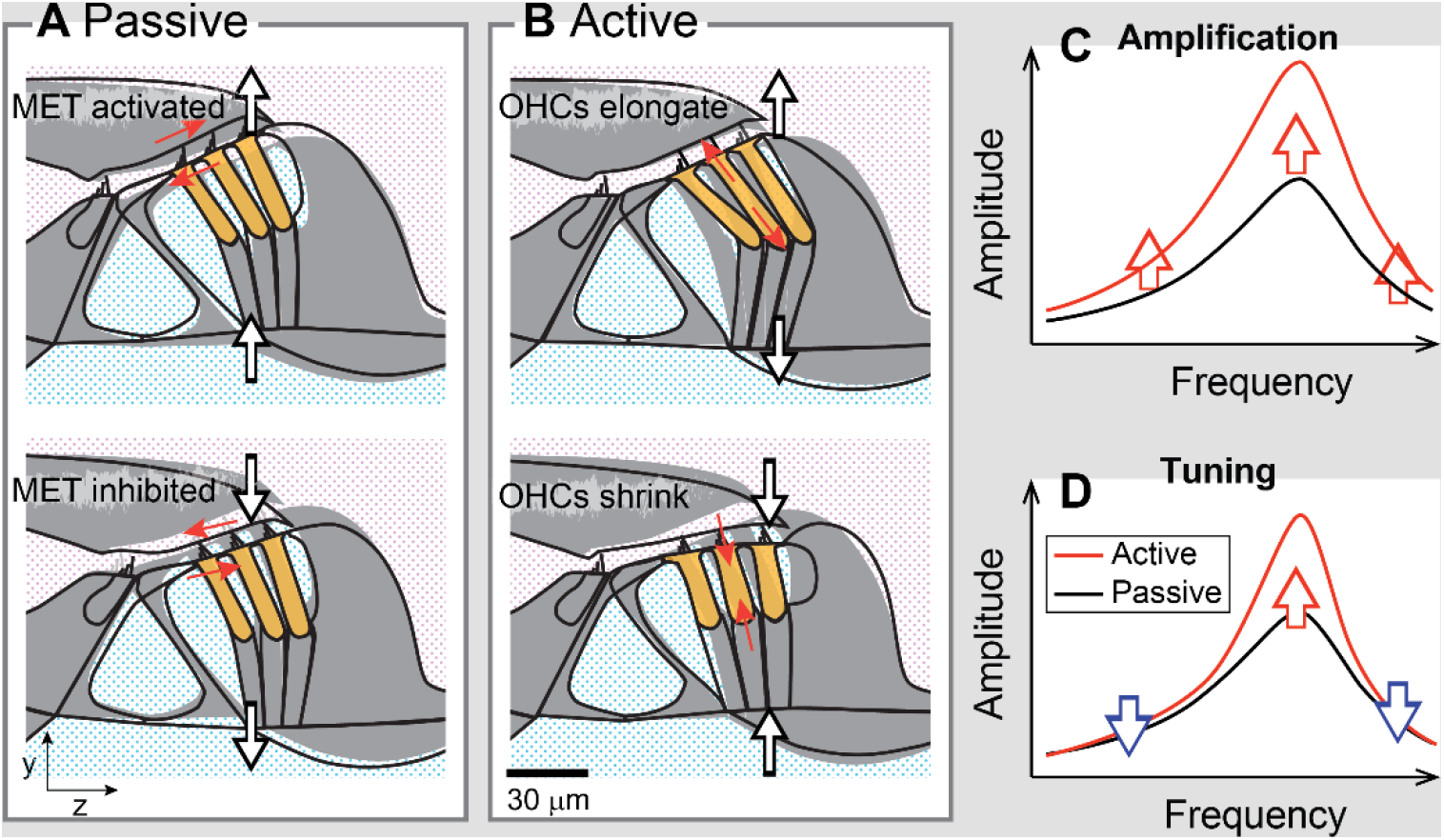
Consequence of destructive interaction. (**A**) Passive vibrations. The OoC vibrates in phase. The relative motion between the tectorial membrane and the reticular lamina results in hair bundle deflection, which activates mechno-transduction. (**B**) Active vibrations. Due to the change in transmembrane potential, the outer hair cells elongate or shrink. When the outer hair cells are motile, fine structures of the OoC vibrate out of phase. (**C**) When active and passive vibrations interact constructively regardless of frequency, the responses are amplified, but tuning quality remains similar. (**D**) Tuning can be enhanced if destructive interaction occurs away from the CF.

In this study, we varied the timing of outer hair cell actuation to examine the idea of destructive interaction between two vibration patterns (Figs. 6 and 7). Before testing the interactions, we characterized the two vibration patterns independently to determine how the OoC vibrates when it is mechanically or electrically stimulated. When mechanically stimulated (passive, Fig. 8A), the vibration pattern was in line with the classical understanding of OoC kinematics. That is, the entire OoC vibrated in phase, and the maximum displacement occurred near the location of the outer hair cells and Deiters’ cells (Fig. 3A). Relative motion between the top and the bottom surface of OoC remained the same over the tested frequency range (Fig. 2). In contrast, when electrically stimulated (active, Fig. 8B), OoC micro-structures vibrated out-of-phase (Fig. 3B). Our observed electrical responses compare better with previous measurements with sensitive (active) cochleae that showed the phase difference between the top and the bottom sides of the OoC (15, 18, 24). In addition to phase variation along the depth of the OoC, our results show apparent phase variation along the radial direction (Fig. 3B). We demonstrated that, depending on the timing (phase) of actuations, there can be either constructive or destructive interactions between the two vibration modes (Fig. 7). If the actuation timing varies over the frequency, there can be constructive or destructive interactions depending on stimulating frequency. Such constructive or destructive interaction could enhance cochlear tuning (Fig. 8D).

It should be reminded that this study is focused on micromechanical aspects, as opposed to macromechanics such as traveling waves. For example, we did not observe a clear sign of traveling waves or tonotopy over the exposed section of cochlear coil. The transfer function in Fig. 7A may reflect artifacts such as wide opening of the scalae and micro-chamber geometry. Physiological characteristic frequency of the tested location is 1.5 kHz, which is different from our nominal characteristic frequency of 0.8 kHz. As a result, our approach cannot be used to estimate physiological tuning quality. With that said, we showed an example of destructive interaction under controlled conditions. To better observe the interaction between vibrations due to different stimulation types, stimulation levels were chosen so that two types of stimulation resulted in similar vibration amplitude. This condition may represent the situations when there is minimal amplification. That is, the results in Fig. 7 may be better compared with the responses in the tail region of traveling waves in natural cochlea.

The destructive interaction may occur under physiological conditions, considering that the timing of outer hair cell actuation varies over frequency. For example, the phase between the reticular lamina and the basilar membrane varies over frequency (15, 16, 36). Ren and his colleagues have discussed the idea of constructive interaction based on the phase difference between the reticular lamina and the basilar membrane (18). Our measurements (Fig. 5) are in line with their view—“*in-phase vibrations of the reticular lamina and basilar membrane result in constructive interference*”. Dong and Olson (37) showed that the phase between the extracellular potential near the outer hair cells with respect to the basilar membrane motion varies over frequency exceeding 90 degrees. It was argued that the phase shift over 90 degrees is an indication of selective amplification (38). Our argument of tuning shares similar reasoning as Olson and her colleagues in that outer hair cell motility away from the best-responding location does not amplify.

New experimental attempts were made in this study to better observe OoC mechanics. It may be worthy to discuss the opportunities and challenges of our approach. First, the micro-chamber was designed to stimulate isolated cochlear tissue both acoustically and electrically. Previous studies using micro-chambers were referred to, including Chan and Hudspeth (39), and Karavitaki and Mountain (22, 30). Besides adopting the merits of those studies (endolymph-perilymph separation, and calibrated electrical/mechanical stimulation), simultaneous application of mechanical and electrical stimulation was newly attempted in this study. This approach was instrumental in isolating and synthesizing different vibration modes within the OoC. Second, we measured individual outer hair cell deformation *in situ* using OCT. Experiments with isolated outer hair cells have been providing biophysical information such as the electro-motile properties of the outer hair cell membrane (40–43). The OoC vibrations due to outer hair cell motility have been providing insights on how outer hair cells contribute to cochlear amplification (32, 39, 44). To our knowledge, the change in the length of individual outer hair cells *in situ* has not been measured before. This was achieved by combining OCT imaging with our microchamber preparation. Compared to the laser interferometry that has long been the gold standard in cochlear vibration measurements, OCT was shown to be efficient in scanning vibrations over a cross-section or a volume of cochlear tissue (19, 24). We achieved an enhanced spatial resolution as compared to existing studies thanks to the removal of bones along the optical path and incorporation of a higher numerical aperture objective (N.A. of 0.4).

There are aspects that have not been fully exploited regarding our experimental approach. First, we did not take advantage of the electro-chemical separation across the OoC. Although we have tested both perilymph-perilymph and endolymph-perilymph conditions, we are not ready to discuss the effects of different chemical conditions, primarily due to the lack of acquired data. The change from perilymph to endolymph in the top fluid space was usually done within 30 minutes following aggressive surgical procedures including the removal of cochlear bones and the cleaving of the stria vascularis. Therefore, it was difficult to tell if any change in the early state was due to the tissue’s settling from surgical procedures, or it was due to the replacement of the top fluid. Besides the effect of endolymph-perilymph, we could have performed different chemical assays such as different calcium levels or the application of mechanotransduction channel blockers, but they have not been examined yet. In the present study, we dedicated a 2-3 hours experimental window to explore OoC mechanics—by measuring OoC vibration patterns due to different types of stimulation. The other aspect that will be helpful but was not implemented in this work is vibration measurement in different directions. For example, the vibrations were measured along one optical axis. We chose the direction approximately parallel to the length of the outer hair cell. As a result, the basilar membrane was tilted about 30 degrees from the optical plane. Had we measured at different orientations of the preparation, we could have distinguished radial and transverse vibrations as was done by Lee *et al*. (36). The radial and longitudinal motions at the base of the outer hair cell were shown to be significant when the OoC was electrically stimulated (22, 30). This study presents the displacement along the axis of the outer hair cell, which is closer to the transverse rather than radial axis.

## Author Contributions

TJ, JCH, and JHN conceived the study. TJ, JB and JHN performed experiments. JHN drafted the MS and the figures. TJ, JHN and JCH edited the MS. JB calibrated experimental devices including the OCT system.

## Acknowledgements

This work was supported by NIH R01 DC014685.

